# Time-resolved transcriptomic of single *V. vinifera* fruits: membrane transports as switches of the double sigmoidal growth

**DOI:** 10.1101/2024.09.27.615328

**Authors:** Stefania Savoi, Mengyao Shi, Gautier Sarah, Audrey Weber, Laurent Torregrosa, Charles Romieu

## Abstract

By revealing that the grape berry loses one H^+^ per accumulated sucrose at the inception of ripening, adopting a single fruit paradigm elucidates the fundamentals of the malate-sugar nexus, previously obscured by asynchrony in population-based models of ripening. More broadly, the development of the individual fruit was revisited from scratch to capture the simultaneous changes in gene expression and metabolic fluxes in a kinetically relevant way from flowering to overripening. Dynamics in water, tartrate, malate, hexoses, and K^+^ fluxes obtained by combining individual single fruit growth and concentration data allowed to define eleven sub-phases in fruit development, which distributed on a rigorous curve in RNAseq PCA. WGCNA achieved unprecedented time resolutions in exploring transcript level-metabolic rate associations. A comprehensive set of membrane transporters was found specifically expressed during the first growth phase related to vacuolar over-acidification. Unlike in slightly more acidic citrus, H^+^ V-PPase transcripts were predominantly expressed, followed by V-ATPase and PH5, clarifying the thermodynamic limit beyond which replacement by the PH1/PH5 complex turns compulsory. Puzzlingly, *bona fide* ALMT kept a low profile at this stage, possibly replaced by a predominating uncharacterized anion channel. Then, the switch role of HT6 in sugar accumulation was confirmed, electroneutralized by malate vacuolar leakage and H^+^ pumps activation.

**Highlights:** To alleviate asynchronicity biases, transcripts showing strict coincidental timing with pericarp physiological phases were disentangled on single berries, enlightening the tight multifaceted membrane developmental control of sugar and acid fluxes.

## Introduction

Developing an edible pericarp to promote the dispersal of mature seeds displays convergent features in Angiosperms. Before coloring and accumulating quick carbohydrates during ripening to attract and provide frugivores an energy reward, green fleshy acidic fruits are protected by astringent proanthocyanidins and undergo an initial expansion period relying on the storage of organic acids as major osmoticums in their hypertrophied vacuoles (Batista-Silva *et al*., 2018; Rauf *et al*., 2019). Whenever as indicated by its name, the major accumulated organic acids may change within species, a similar acid trap mechanism operates, based on keeping the vacuolar pH (pHv) below the pK of the organic acid so that Henderson-Hasselbalch equation forces transported anions to recombine with H^+^ in the vacuolar lumen, thereby alleviating membrane depolarization, reducing pHv decrease, and subsequent transport retro inhibition (Martinoia *et al*., 2007). This obvious lack of charge neutralization by K^+^ also prevents the depletion of this mineral in the rhizosphere of a perennial plant amenable to decades of fruit dissemination. Nevertheless, reaching an extreme pHv raises a bioenergetic challenge that is solved by recruiting a low stoichiometric ratio P-type ATPase complex in lemon varieties with a pHv of about 2.5 (Strazzer *et al*., 2019). Not only should H^+^ V-ATPase and H^+^ V-PPase be unable to reach such pHv beyond their expected thermodynamic limits, but they would even collapse it by functioning in the reverse mode, which is prevented by the unique conformation of the inhibitory H subunit in citrus V-ATPase (Tan *et al*., 2022). The predominant PH1/PH5 complex in citrus is transactivated by the basic helix-loop-helix (bHLH) transcription factor *Noemi* (Butelli *et al*., 2019), which in grapevine controls the possibly unrelated processes of acids and proanthocyanidins accumulation upon acting in an MYB/bHLH/WD40 (MBW) complex, whose orthologs in grapevine are represented by the bHLH *VviMYC1* (Hichri *et al*., 2010), the TFs *VviMYB5b* and *VviWRKY26* (Amato *et al*., 2019). The PH5 ortholog is apparently expressed in grapevine, but unlike in citrus, proteomic, enzymatic, or vectorial evidence show that functional V-ATPase and V-PPase largely predominate in green grape berries at pHv 2.7 (Terrier *et al*., 1998; Rienth *et al*., 2016; Kuang *et al*., 2019). Furthermore, aluminum-activated malate transporters (ALMT), such as the apple *ALMT9*, or *Ma1* gene (Bai *et al*., 2012), play a critical role in fruit acidification upon transferring malate inside the vacuole. Surprisingly, despite displaying suitable inward rectifying transport of malate and tartrate, the gene described as *VviALMT9,* is preferentially expressed during the ripening period rather than during the acid accumulation phase (De Angeli *et al*., 2013), which questions its practical implication in building grape acidity. A better temporary resolution would shed light on a situation that is all the more puzzling as the quite uncommon tartaric acid, stronger than malic acid, accumulates simultaneously with proanthocyanidins in the very young berry. It is becoming increasingly clear that such energy accumulated as an H^+^ gradient will play a key role during ripening. Critical re-evaluations of sugar accumulation versus malate breakdown at this stage led to observing an initial 2 hexoses/H^+^ stoichiometry in all *vinifera* varieties investigated so far (Shahood *et al*., 2020; Bigard *et al*., 2022), in line with the induction of the *VviHT6* transcript and protein that has been widely confirmed (Kuang *et al*., 2019; Savoi *et al*., 2021) since the initial report of Terrier *et al*. (2005). This raises the question of the exit route of malate before H^+^ pumping back by V-ATPase and V-PPase takes over the electro-neutralization process, as evidenced on tonoplast vesicles (Terrier *et al*., 2001).

Grapevine (*Vitis vinifera*) is a major fruit crop and a model for non-climacteric fruits. The latest process-based modeling studies are rooted in the crude approximation that berry development would reflect the compositional changes of the future harvest as an average population and reciprocally (Zhu *et al*., 2019; Tornielli *et al*., 2023). Individual berry approaches show that current models of berry development are biased by asynchronous ripening within the future harvest (Bigard *et al*., 2019; Shahood *et al*., 2020; Daviet *et al*., 2023). Indeed, asynchronicity lies within the duration of phenological stages under study, making average kinetics developmentally chimeric. Consequently, single-fruit transcriptomics yielded a straightforward identification of genes suddenly switched off with sugar accumulation at the end of ripening and provided decisive insights into the structure and energetics of the phloem unloading pathway (Savoi *et al*., 2021). Such a proof of concept challenges us to revisit the whole berry developmental cycle from the strict kinetic and quantitative point of view.

Recently, several studies have been dedicated to deciphering the signaling cascade triggering the onset of ripening, a pivotal developmental point in fruit physiology. A quite consensual hormonal pattern emerged, but which family or transcription factor members are responsible for the transcriptomics reprogramming is still being determined. In non-climacteric fruits, a slight increase of endogenous ethylene hormone is followed by a larger peak of abscisic acid onsetting the veraison, prevented by auxins (as reviewed in Perotti *et al*., 2023; Zenoni *et al*., 2023). On the other hand, a whole range of NACs (D’Incà *et al*., 2023), GRAS (Neves *et al*., 2023), bHLH, WRKY (Fasoli *et al*., 2018), LOB (Grimplet *et al*., 2017) transcription factor families and two ripening time control (RTC1 and RTC2) proteins (Theine *et al*., 2021) would trigger ripening molecular events, without defining a conclusive ripening cascade. The TF *VviERF27* was recently identified as the single candidate in the veraison locus *Ver1* (Frenzke *et al*., 2024), influencing ripening start. In this respect, results from these disparate studies need to be synchronized on a common time frame relying on unambiguous samples and internal clock markers.

Here, we report the first single fruit transcriptome study encompassing the complete three-month period from anthesis to over-ripening in the world-cultivated Syrah variety. This new approach yielded a straightforward identification of key transporters and bioenergetic players switching abrupt metabolic changes during pericarp development. Ultimately, we attempted to individuate the expression profiles of pivotal master regulators of the berry ripening process.

## Materials and Methods

### Plants

The *Vitis vinifera* variety Syrah, an iconic variety for the production of red wines in regions under temperate climates, was considered for this study in 2018 and 2019. The vines, grafted on SO4 and planted in 2000 in the experimental facility of Institut Agro Montpellier, were irrigated daily during the summer to avoid severe water shortage. Classical phytosanitary treatments were applied over the season to ensure healthy plant and fruit development.

### Single berry sampling

The double sigmoidal growth pattern of single berries was monitored through recurrent pictures of selected clusters on different plants, starting at flowering and continuing until two weeks after maximum berry volume. Photographs were taken using a Lumix FZ100 (Panasonic), keeping the focal range and cluster to camera lens distance (30cm) constant. The volumetric growth of single berries was calculated following picture analysis with the ImageJ software. After calibrating all images with their 1 cm internal standard, berry edges were manually adjusted to ellipsoids whose areas were determined by pixel counts before estimating the berry volume as previously described (Savoi *et al*., 2021).

The complete developmental period was followed for twenty berries, allowing to model their individual double sigmoidal growth patterns, while a second set of 219 berries were sampled according to their own volume changes over two seasons (2018 and 2019) before sugar, organic acids, and K^+^ were analyzed on circa 0.1 g aliquots of frozen samples, as described below. Calendar dates were converted to individual DAF ones, following resynchronization based on berry growth, composition, and softening dates. The net fluxes of water (growth), organic acids, and sugar accumulation were calculated for each berry before selecting the eleven time points for RNAseq samples (Table S1). To avoid circadian cycle influences (Rienth *et al*., 2014; Davies *et al*., 2023), pictures and berries were sampled at the same time of the day, between 9 and 11 AM. Berries without pedicel were rapidly deseeded before freezing in liquid N_2_ (within 1 min after harvest) and stored at -80°C. A total of 219 single berries were ground to a fine powder under liquid N_2_ using a stainless steel ball mill (Retsch MM400).

### Primary metabolites analysis

Each single berry was analyzed for sugars (glucose and fructose) and acids (malic and tartaric acids) by high-performance liquid chromatography. For each 219 berries, 100 mg of frozen powder was diluted 6x with a solution of HCl 0.25 N, well-shacked, and left overnight at room temperature. Samples were then centrifuged at 13,000 g for 10 min, and a supernatant aliquot was diluted 10x with a solution of H_2_SO_4_ 5mM containing 600 µM of acetic acid as internal standard. Samples were transferred to HPLC vials and injected, according to Rienth *et al*. (2016). For K^+^ analysis, the previous samples in HCl 0.25 N were further 10x diluted with MilliQ water and analyzed with atomic absorption spectroscopy as in Bigard *et al*. (2020). Data were expressed as mEq for organic acids or mM for sugar concentration; content per fruit (concentration x volume) was defined as mEq/Nfruit or mM/Nfruit where N represents the number of fruits needed to reach 1 kg FW at maximal volume, which was 485 for Syrah.

### RNA extraction and sequencing

To resynchronize berries at homogeneous developmental stages, single berry triplicates (pericarp) were selected based on relative growth, sugars, and organic acids, yielding 33 samples before individual RNA extraction and library preparation as in Rienth *et al*. (2016). Samples were sequenced on an NGC Illumina HiSeq3000 in paired-end mode, 2×150 bp reads, at the Genotoul platform of INRAe-Toulouse.

### Transcriptomics data analysis

Raw reads were trimmed for quality and length with “Trimmomatic”, version 0.38 (Bolger *et al*., 2014). Reads were aligned against the reference grapevine genome PN40024 12X2 (Canaguier *et al*., 2017) using the software “Hisat2”, version 2.1.0 (Kim *et al*., 2015), yielding an average of 31.6 M sequences per sample. Aligned reads were counted with “HTSeq-count” (version 0.9.1 (Anders *et al*., 2015)) using the VCost.v3 annotation. Genes were filtered by applying an RPKM>10 cut-off in at least one experimental condition, resulting in 7266 expressed genes. Transcripts expressed as normalized RPKM were tested for multi-time-series significance to screen those genes showing significant temporal expression changes using the “MaSigPro” R package (Nueda *et al*., 2014) with parameters degree=7, rsq=0.7. The output, consisting of a list of 6374 time-modulated genes, accompanied by osmolytes metabolic rates and phenological stages, was subjected to a weighted correlation gene co-expression network analysis with the “WGCNA” R package (Langfelder and Horvath, 2008) using the RPKM dataset, with parameters ß power 30, network type signed, min module size 100, deepSplit 1, MEDissThreshold 0.1. WGCNA resulted in the 13 following modules: black (560 genes), blue (660), brown (793), green (952), yellowgreen (296), lightcyan (103), magenta (648), midnightblue (113), pink (348), tan (266), turquoise (693), yellow (560), and finally grey (382) containing genes unassigned.

### Calculation of the transcriptomics clock

A PCA was conducted on unfiltered variance stabilized transformed (VST, DESEQ) gene expression data from the 33 sequenced samples. The transcriptomic time was calculated as the curvilinear distance between the orthogonal projections of successive individual berry data on a fitted polynomial line in the first PCA plane. The curvilinear distance was approximated as the integral of dl = R(φ).dφ following a change from cartesian to polar coordinates. The rate of accumulation or degradation for each osmolyte was then retrieved based on the transcriptomic time and the metabolic data.

## Results

### Single fruit monitoring provides new kinetic bases for berry development

Twenty individual berries were photographed every two to three days for three months, starting a few days after flowering (DAF). Image analysis led to characterize typical double sigmoidal growth curves followed by two weeks of berry shriveling, as accepted in the grapevine, except for the facts that the herbaceous plateau appeared particularly long with a continuous slow berry volume increment and the second intense growth period lasted 3.5 weeks, as previously highlighted on individual berries (Shahood *et al*., 2020). Best fits with the 4-parameter sigmoidal function v=v_0_+v_max_/(1+exp((t_0_-t)/b)) were obtained for each growth phase, where v stands for fruit volume, t for time (days), v_0_ and v_max_ initial volume and plateau volume, t_0_ the time at which both mid-growth and maximal growth rate were reached (Fig. 1A and Table S2, S3). Asynchronicity was largely eliminated upon resynchronizing t_0_ values, shifting from calendar time to single fruit-specific DAF. Average values of 23.14 and 62.05 DAF were obtained for t_0_ green and t_0_ ripening (i.e., for mid-first and mid-second intense growth phases, respectively), resulting in an offset of 39 days. Following the normalization of individual berry peak volumes, the first (or green) growth period plateaued at 0.44 (Fig. 1A, Table S3), indicating a greater expansion during the second growth period (ripening). Moreover, according to their respective maximal growth rates, the ripening berry expands 1.74 times faster than during the green stage. Terminal shriveling was more or less pronounced on different berries. On average, the berry volume decreased by 17% in three weeks after the stop of phloem unloading, with a minimum decrease of 7% and a maximum of 29% (Fig. 1A, Table S2). This decrease was not correlated to the berry surface-to-volume ratio (not shown). On these 20 berries and those destructively sampled, the first intense growth phase lasted until 35 ± 1.5 DAF on average, and softening (onset of ripening) occurred at 51 ± 1.5 DAF. Berry skin coloration followed softening by about one week, starting at about 58 ± 1.5 DAF, quite simultaneously with the second intense growth phase. Maximum berry volume was reached at 75 ± 1.5 DAF, 24 days after softening (Table S1).

**Figure 1.**
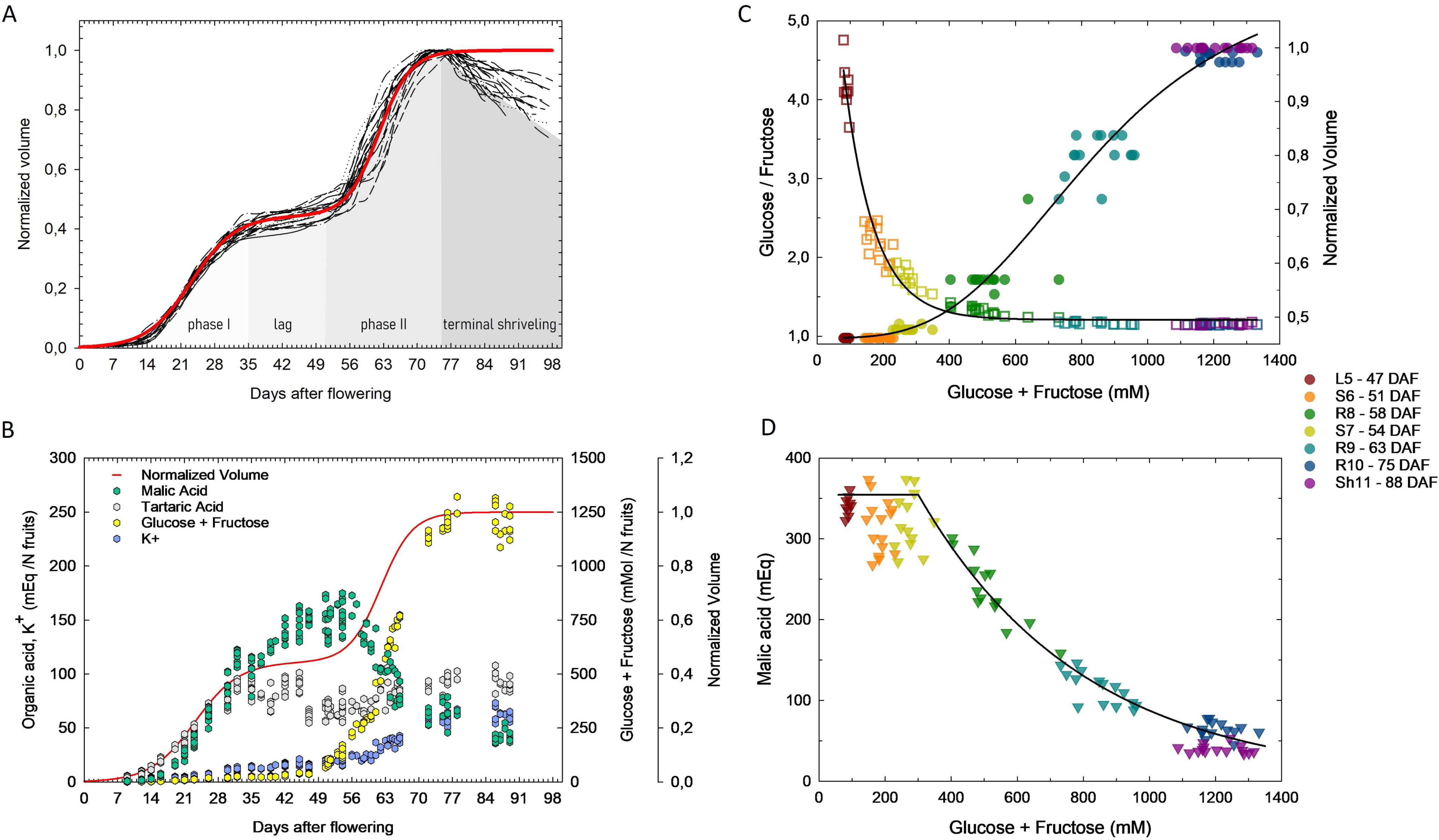
Growth and major osmotica in synchronized berries. **A**: Growth kinetics of 20 single Syrah berries (black lines). The maximal volumes were normalized, and then the first growth periods were synchronized upon shifting the individual DAF. Experimental data (black lines) were fitted to a double logistic function (red line): v=0.44506/(1+exp((23.14-time)/4.78848)) + 0.55494/(1+exp((62.05-time)/3.31967)), Table S3. The berry developmental phases are named according to the different shades of gray. **B**: Growth kinetics and accumulation of major osmotica in synchronized berries, from flowering to over-ripening. For comparative purposes, units relate to N=485 berries, yielding 1 Kg of fruit for variety Syrah at maximum volume. Green and gray dots represent malic and tartaric acids, while yellow and blue represent glucose+fructose and potassium, respectively. The detailed fluxes dataset is available in Table S4. The red line reports the fitted double sigmoidal growth curve established in Fig. 1A. **C**: Sugar accumulation vs. the ratio of the two hexoses. Growth resumption occurs later (at 0.4M) than softening and the onset of sugar accumulation. Squares represent the evolution of the glucose/fructose ratio, while circles show berry expansion. **D**: Sugar accumulation vs. malate breakdown. L5 green hard, S6-S7 green soft, R8-R9-R10-Sh11 red to black. At Sh11, the decrease in berry volume was not accompanied by visible berry wrinkling yet.

### Acids storage and breakdown during fruit development

Tartaric acid, already present at 5 mEq/Nfruits at 9 DAF (Table S4), the first sampling date, reached circa 87 mEq/Nfruits at 35 DAF (Fig. 1B) and remained stable after that, regardless of a slight inflection at 47 DAF (the first sampling date of the second experimental season), possibly due to different environmental conditions during the first growth phase (Becker *et al*., 2022; Reluy *et al*., 2022), even if data from Melino *et al*. (2009) showed the same inflection at softening in a non-synchronized population of berries. Tartaric acid was still at 91 mEq/Nfruits at peak volume (75 DAF), which confirms it is only accumulated during the first growth phase and not further metabolized during ripening. By contrast, malic acid, being only 0.7 mEq/Nfruits at 9 DAF (Table S4), started to accumulate a few days later than tartaric acid but continued until reaching a malate/tartrate ratio above 2 (151 mEq/Nfruits) at circa 47 DAF (Fig. 1B). Malic acid was roughly constant around softening (51 DAF) and suddenly dropped at about 58 DAF, displaying a pseudo-first-order decrease (Fig. 1D) until reaching values of 41 mEq/Nfruits at 88 DAF. Our data distinctly states that the organic acid present in the berry pericarp at our first metabolic sampling was tartrate, which anticipated malate accumulation by a few days (Fig. 1B). Then, malate became 2-3 fold higher than tartrate during the first growth period before ripening was triggered. The net accumulation rates of tartaric and malic acid, the major osmoticums in the first water accumulation period, were estimated for the first time on a per-fruit basis, exhibiting respective values of 3 and 4.4 µEq/min.NFruits when reaching a quasi-constant accumulation rate, as can be seen in Fig. 1B.

Potassium slowly accumulated during the green and lag stages (from 1 to 16 mEq/Nfruits), accelerated during the ripening phase before reaching 59 mEq/Nfruits (Fig. 1B, Table S4) at phloem arrest (Savoi *et al*., 2021). Puzzlingly, K^+^ accumulation was relatively insignificant during the malate breakdown period accelerating afterward (Fig. S1).

During the green stage, the elevated glucose/fructose ratio above 3 (Fig. 1C) illustrates the preferential use of fructose from imported sucrose as a carbon and energy source. However, fructose consumed in excess of glucose should just allow the synthesis of 60% accumulated malic acid, implying that a supplementary part of imported sucrose is entirely metabolized as organic acids or pyruvate as a respiratory substrate. Actually, photosynthesis would never reach the compensation point in berries (Niimi and Torikata, 1979).

### The sequence of events at the onset of ripening and sugar accumulation

Sequential destructive analysis of single berries confirmed that the first phenotyping signs of ripening were simultaneous softening and decrease of the G/F ratio below 3 (Fig. 1C), subsequent to the marked acceleration of sucrose import and hydrolysis above 0.15 M hexoses, while berry expansion resumed later, at 0.4 M hexoses. The combination of growth and concentration data showed that at softening (51 DAF), glucose and fructose accumulation progressively accelerated up to 63.7 µmol hexose.min-1.Nberry-1, representing a 20-fold acceleration compared to the green stage (3.1 µmol hexose.min-1.Nberry-1), before abruptly stopping 24 days later. However, the initiation of malic acid breakdown did not occur before 0.3 M sugar (Fig. 1D). Once induced, the initial rate of malate breakdown was in line with a 1 sucrose/1 H^+^ exchange at the tonoplast (Fig. S1). Finally, it is important to note that K^+^ accumulated at a 25x lower rate than sugars during the ripening phase.

### Transcriptomic overview

For RNA sequencing, thirty-three berries representing eleven sub-phases of berry development were selected among the 219 sampled ones, according to their individual growth rate and primary metabolite contents. The resynchronized DAF of the triplicates of single berry and their developmental and phenological acronyms are detailed in Table S1 and visualized in Fig. 2A. 54% and 15% of the variance in gene expression were distributed on the two first axes of the principal component analysis (PCA) (Fig. 2B). The three single fruits remained largely indistinguishable inside biological replicates, even if slight divergences appeared at the R9 stage and beyond. The developmental stage clearly drove sample distribution. Berries did not depart from the same line, illustrating the higher accuracy of the transcriptomic clock compared to multi-trait phenotyping to synchronize berries. As much as 14% of the top 100 influencing genes driving sample distribution on the negative side of PC1 (from G1 to L5) encoded photosynthesis-related functions (Table S5). Noticeably, one-half of the 100 most influencing transcripts on the PC1 positive side (from S6 to Sh11) have been previously identified as “switch” genes (Palumbo *et al*., 2014) transcriptionally induced at softening and significantly activated during ripening. Therefore, PC1 sample separation confirms the general transcriptomics reprogramming during the ripening process (Fasoli *et al*., 2012), clearly separating the first growth phase (including the green plateau) from the ripening second one. Interestingly, the genes placing samples in the negative section of PC2 (i.e., G1, G2, partially R9, R10, and Sh11) (Table S5) were more particularly devoted to the phenylpropanoid and flavonoid pathways (but not anthocyanin specific, which were listed in PC1+). By contrast, positive gene loadings for PC2 did not belong to a particular class or known pathway.

**Figure 2.**
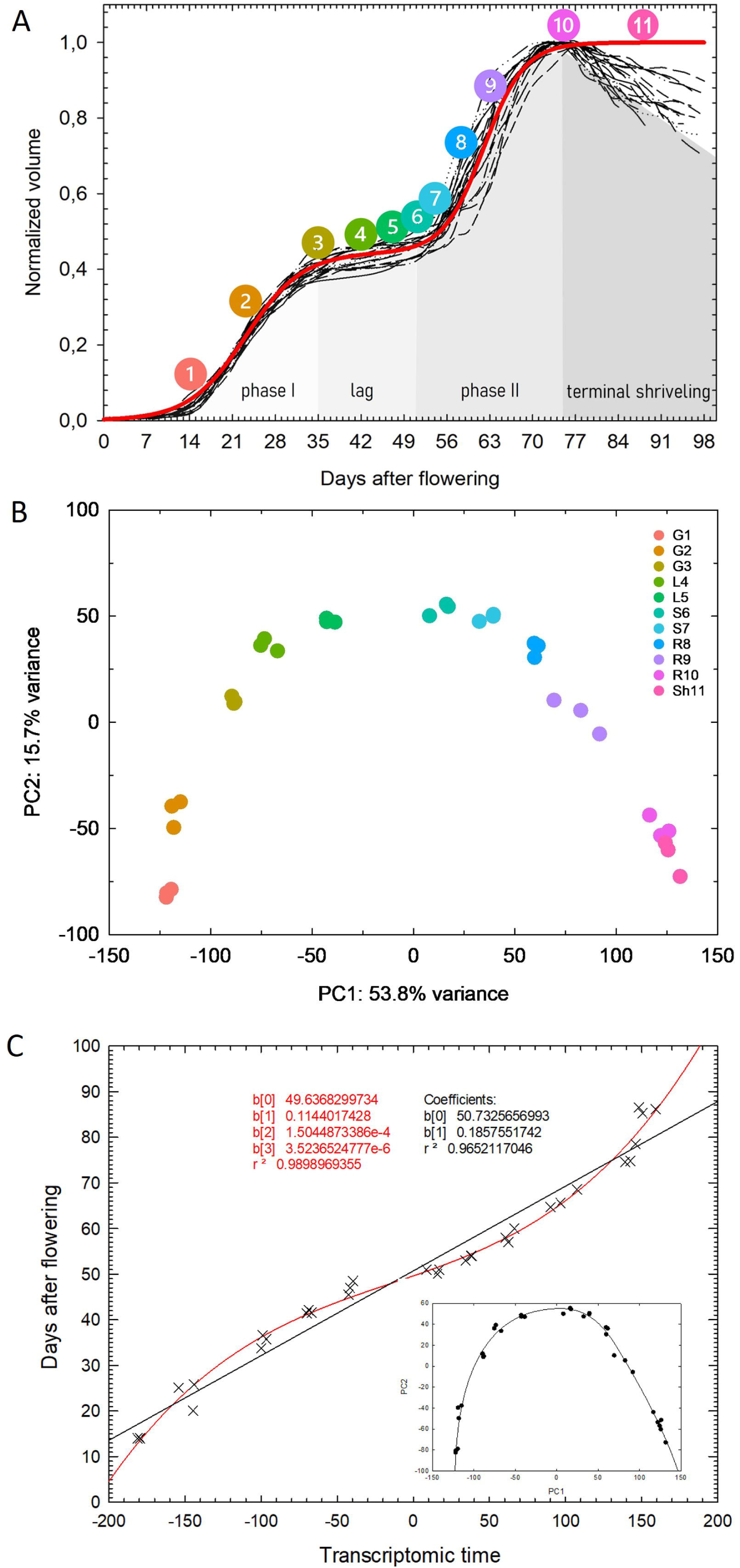
Transcriptomics clock overview. **A:** Repartition of the eleven developmental stages subjected to RNA-Sequencing according to berry growth phases. **B:** PCA of gene expression in 33 single berries sampled at 11 developmental stages, as color legend specifies. G: green phase, L: lag phase (green hard), S: softening (green soft), R: ripening red-purple, and Sh: shriveling. Green phase: G1 to L5, Ripening: S6 to R10, Over Ripening: Sh11. **C**: Transcriptomics time (red line, Table S6) is inferred from the curvilinear distance of the fitted line on the PCA (inlet).

### The single-berry transcriptomic clock

The curvilinear distance on the fitted line passing through the 2D PCA of single-berry transcriptomes was found to be roughly proportional to their resynchronized internal time, which still suffers from uncertainties owing to the complexity of berry resynchronization based on individual growth curves and primary metabolites (Fig. 2C). A significant acceleration of transcriptomic changes occurred around softening, while homeostasis was reached at maximum volume or phloem arrest. On individual berries, this approach, conceptually inherited from Tornielli *et al*. (2023), virtually eliminated the random noise on uncharacterized, thus variable mixes of unsynchronized berries. Indeed, single berry transcriptomic time (Table S6) obtained blindly from unfiltered RNAseq results allows dating individual berries with a precision within the few days range, if not less, and outperforms all alternative resynchronization procedures, as illustrated on post-veraison samples.

### Weighed gene co-expression network analysis

After filtering for low-expressed genes, 6374 genes were significantly modulated over time. Their subdivision into 12 different modules of clusters of highly correlated genes (Fig. 3A, 3B, Table S7) (disregarding the grey module that includes all unassigned profiles) allowed us to study the transcriptomic regulation of metabolic fluxes and related traits with unprecedented time precision.

**Figure 3.**
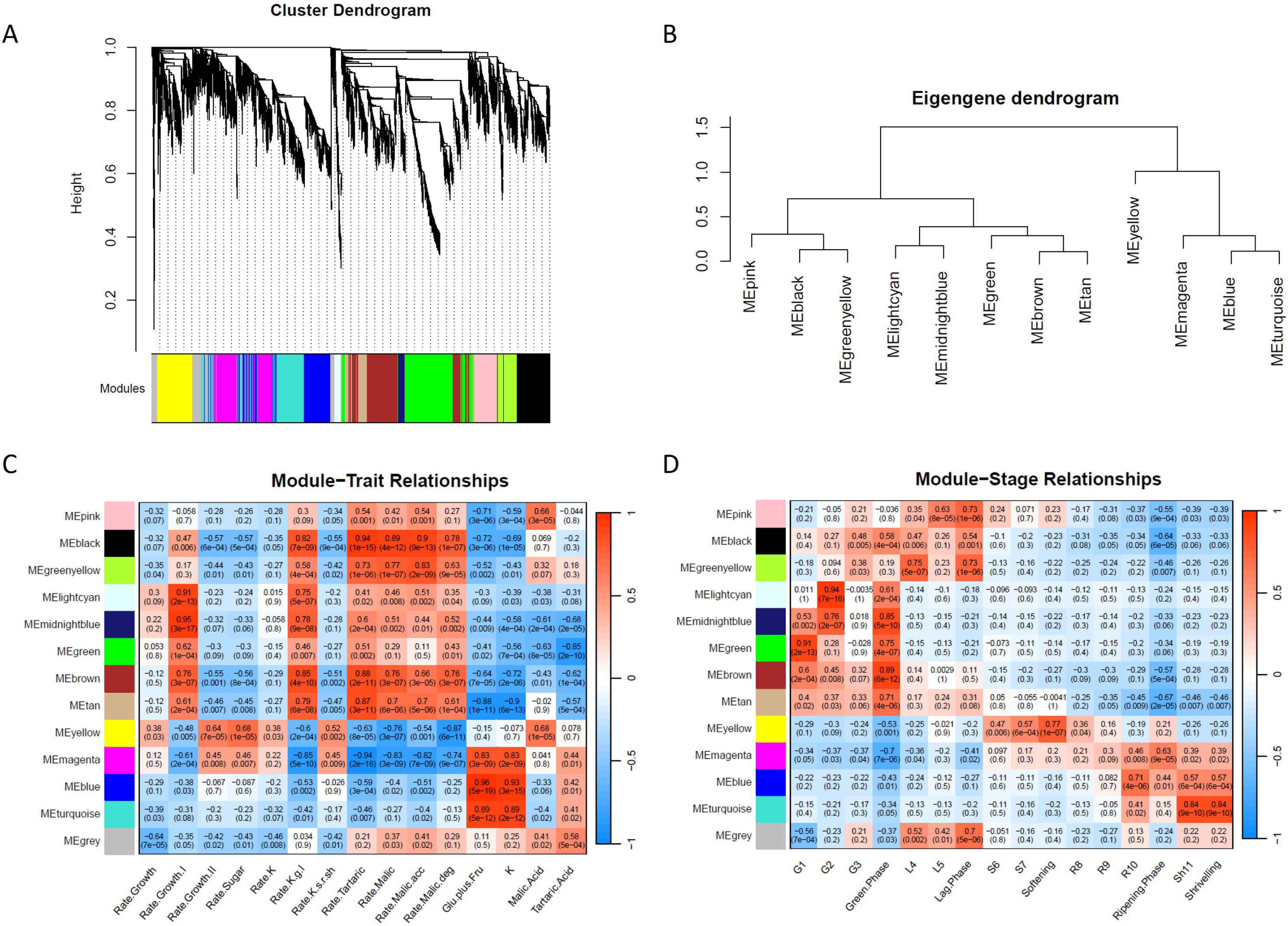
Weighted Gene Coexpression Analysis results. **A:** Hierarchical clustering tree dendrogram of 6374 time-modulated genes, with dissimilarity calculated with a soft thresholding power β of 30 based on topological overlap matrix. Each leaf (black vertical lines) corresponds to a gene. Branches of the dendrogram group together densely interconnected, highly co-expressed genes assigned 13 module colors, visible on the bottom bar. **B:** The Eigengene dendrogram shows the modules’ distribution and interrelationships. ME = Module Eigengene. **C and D:** Module-Trait relationships and Module-Stage relationships, respectively. Each row corresponds to a module, and each column corresponds to a trait specified at the bottom. Numbers inside the cell describe the correlation coefficient and the associated p-value in brackets. Red and blue colors indicate a positive or negative correlation between the trait or stage and the module. Modules are listed on the right sidebar column.

No specific gene expression displayed a significant association with the berry growth rate on both phases of the double sigmoidal curve (Fig. 3C). Examining the two growth phases separately led to the identification of the lightcyan and midnightblue modules as highly correlated (0.91 and 0.95) with the first growth phase, followed by brown, green, and tan modules (0.76, 0.62, 0.61), (Fig. 3C). Modules black, brown, yellowgreen, and tan correlated with the rate of tartaric acid (0.94, 0.88, 0.73, and 0.87) and malic acid synthesis (0.89, 0.76, 0.77, and 0.70), and the green phase in general, with some differentiation among G1 and G2 (Fig. 3C, 3D). Contrary to the growth rate, separating malic acid accumulation in the green phase from its degradation during ripening, did not improve gene association with the net accumulation rate (either positive or negative) during the complete cycle. The lag phase and then L4 and L5 separately were associated with yellowgreen (0.73 for the lag, 0.75 for L4) and pink (0.73 for the lag and 0.63 for L5) modules. The second growth rate was associated with the yellow module (0.64) and the magenta (0.45) to a lesser extent. Similarly, the yellow module correlated with the accumulation rate of sugar (0.68), softening (0.77), S6 (0.47), and S7 (0.57). None of the modules showed a high correlation with the rate of K^+^ storage. However, when the K^+^ accumulation rate was considered separately during the green-lag and then softening-ripening phases, we could notice that modules brown, black, tan, midnightblue, lightcyan, and greenyellow correlated with the berry first phase of growth behavior, while yellow and magenta correlated with the ripening one. The ripening phase was associated with the magenta module (0.63), while the terminal shriveling was with the blue (0.57, and R10 with 0.71) and turquoise (0.64, and Sh11 with 0.84) ones. At the same time, sugar concentration, but not accumulation rate, was strongly associated with the blue (0.96), turquoise (0.89), and magenta (0.83) modules, as for K^+^ (0.93, 0.89, 0.83).

### Transcriptomic regulation of water fluxes and cell expansion

Different subsets of aquaporins on the plasma or the tonoplast membranes and cell wall-related genes were specifically expressed during each growth phase, both associated with an eightfold expansion of the average cellular volume (Ojeda *et al*., 1999). In particular, the PIPs, TIPs, and EXPs expressed during the green phase were found in the black, tan, brown, and green modules, while the ones activated at softening belonged to the yellow module only; the two EXPs up-regulated later during ripening were grouped in the magenta module instead. Moreover, genes of the same family were expressed at different magnitudes, enlightening different genetic controls over these genes.

Specifically, two PIPs (*VviPIP2.7 - Vitvi03g00155 and VviPIP2.4 - Vitvi06g00281)* and two TIPs (*VviTIP2.1 - Vitvi09g00329 and VviTIP1.1 - Vitvi06g01346*) exhibited an early green stage-specific expression, as majorly emphasized by *VviPIP2.7* and *VviTIP2.1*, thus allowing a continuous influx of water in green expanding berries, while showing a fast decrease during the lag phase (Fig. 4A, 4B). Then, at softening (51 DAF), these ‘green’ isogenes were mainly replaced by the expression of *VviPIP2.5* (*Vitvi13g00605*) and *VviPIP2.3* (*Vitvi08g01038*) on the plasma membrane and by *VviTIP1.2* (*Vitvi08g01602*) and to a lesser extent *VviTIP1.3* (*Vitvi13g00255*) on the tonoplast (Fig. 4A, 4B). Furthermore, *VviPIP1.3* (*Vitvi02g00310*) and *VviPIP1.4* (*Vitvi15g01110*) were expressed during both phases of berry growth until being repressed at the phloem arrest, stopping the influx of water into the berry. *VviEXPA16* (*Vitvi14g01977*) was the most expressed expansin gene during the first growth phase and was slightly enhanced in the second one as well. A rather low RPKM, *VviEXPB02* (*Vitvi12g00342*), was expressed only in the first growth phase. Three other isogenes were abruptly switched on at softening, with *VviEXPA14* (*Vitvi13g00172*) and *VviEXP19* (*Vitvi18g00189*) showing the highest expression at the end of the lag phase; a similar trend was observed for the less expressed *VviEXP18* (*Vitvi17g01251*). These genes were progressively replaced by two other genes (*VviEXPA01* - *Vitvi01g01030* - and *VviEXPB04* - *Vitvi15g00643*) induced during the second growth phase and clearly peaking with berry growth rate at 58 DAF, still substantially expressed during over-ripening, indicating a role in the turn-over of cell wall structure during the shriveling period (Fig. 4C, 4D).

**Figure 4.**
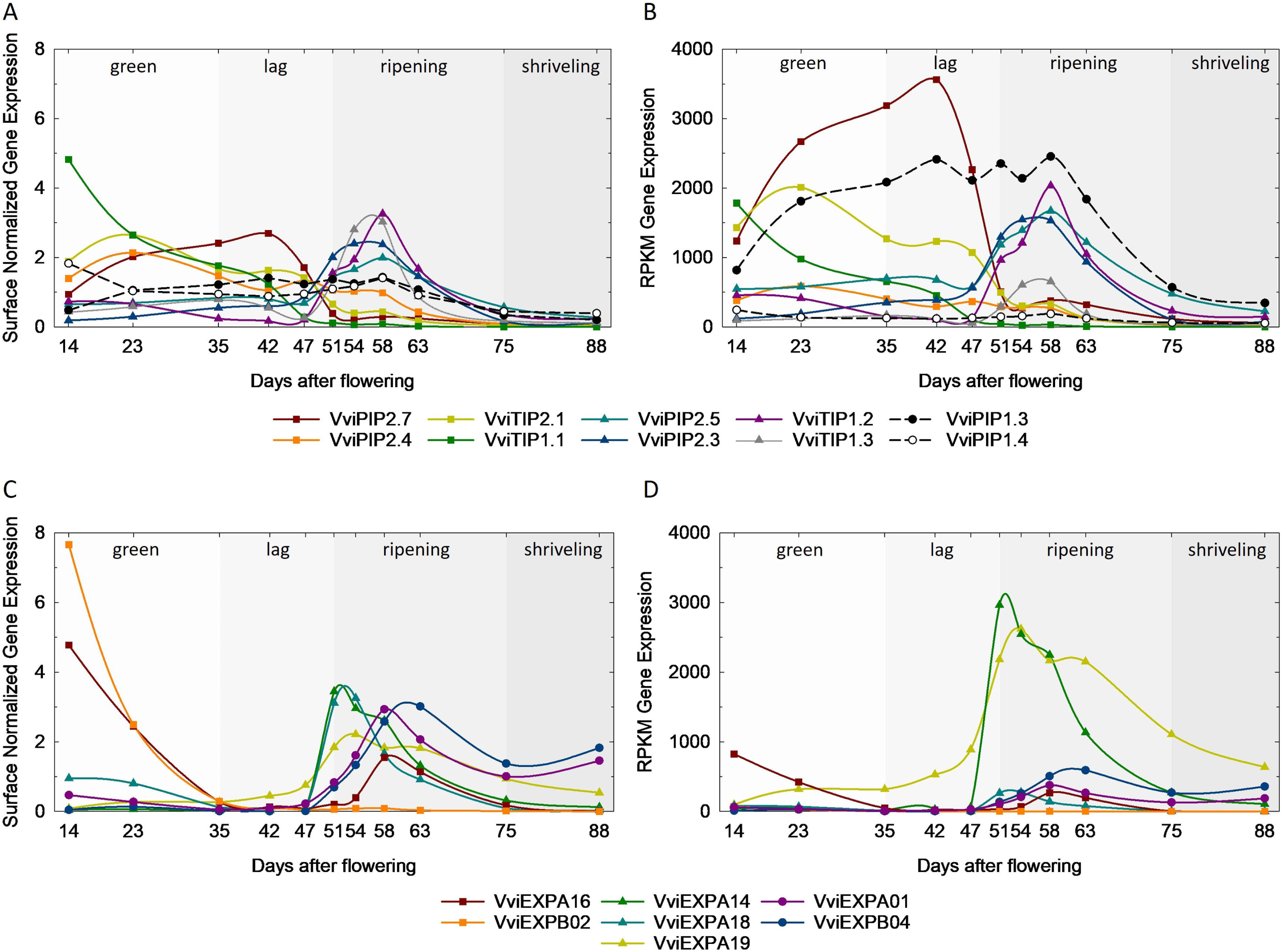
Fruit growth and expansion. Aquaporin (**A, B**) and expansin (**C, D**) genes active in the first period of water entry and growth phase as *VviPIP2.7*, *VviPIP2.4*, *VviTIP2.1*, *VviTIP1.1*, *VviEXPA16* and *VviEXPB02* and the isogenes active in the second phase of growth as *VviPIP2.5*, *VviPIP2.3*, *VviTIP1.2*, *VviTIP1.3*, *VviEXPA14*, *VviEXPA19*, and *VviEXPA18*. In addition, two more expansion genes were identified with an increasing trend during the shriveling time: *VviEXPA01* and *VviEXPB04*. Lastly, two aquaporins were active in both green and ripening phases (*VviPIP1.3* and *VviPIP1.4*). Graphs are expressed as surface normalized data (A, C) and Reads Per Kilobase per Million mapped reads (RPKM) (B, D). In the surface normalized graphs, each expression value was divided by the average on the 11 developmental points to reject large differences in expression among genes. Softening (S) started at an unknown date between 47 and 51 days.

### Proanthocyanidins and organic acid

Five genes committed to the proanthocyanidins biosynthetic pathway or its regulation were specifically expressed at the very beginning of berry development (green module), notably leucoanthocyanidin reductase 1 - *VviLAR1* (*Vitvi01g00234*), anthocyanidin reductase - *VviANR* (*Vitvi10g02185*), the PA transporters *VviPA-MATE1* (*Vitvi12g00101*) and *VviPA-MATE2* (*Vitvi12g00099*), the TF *VviMYBPA2* (*Vitvi11g00099*) (Terrier *et al*., 2009; Pérez-Díaz *et al*., 2014). However, both *VviMYBPA1* (*Vitvi15g00938*) and *LAR2* (*Vitvi17g00371*) were clearly delayed (brown module) (Fig. 5A, 5B). The anthocyanins *VviMYBA1* (*Vitvi02g01019*), *VviMYBA2* (*Vitvi02g01015*), their target gene *VviUFGT* (*Vitvi16g00156*), and the anthocyanin MATE transporter (*Vvi-A-MATE*, *Vitvi16g01913*) showed a simultaneous increase in expression a few days after softening (54 DAF, magenta module). One of the most expressed genes (in RPKM) at 14 DAF was *VviVTC2* (GDP-mannose 3,5-epimerase 1, *Vitvi19g00549*), the regulatory step in the ascorbate pathway in plants, as the precursor for tartrate in grape, which exhibited a green stage-specific expression pattern (Fig. 5C, 5D). The expression of *VviL-IDH3* (L-idonate hydrogenase, *Vitvi16g01858*), the rate-limiting enzyme of tartrate synthesis, was also maximal at 14 DAF, decreasing in the green stage until virtually annihilating during the sugar accumulation process, as in DeBolt *et al*. (2006). Unexpectedly, this gene was reactivated in shriveling berries, contrary to *VviVTC2*. These genes were placed in the brown and green modules, showing a greater correlation with tartaric acid and stage G1. A far less documented, multifunctional enzyme in the late tartaric acid biosynthesis pathway, *Vvi2-KGR* (2-keto-L-gulonic reductase, *Vitvi09g00358*), was also preferentially expressed at 14 DAF before reaching a steady-state level, precisely as in Jia *et al*. (2019) (Fig. 5C, 5D).

**Figure 5.**
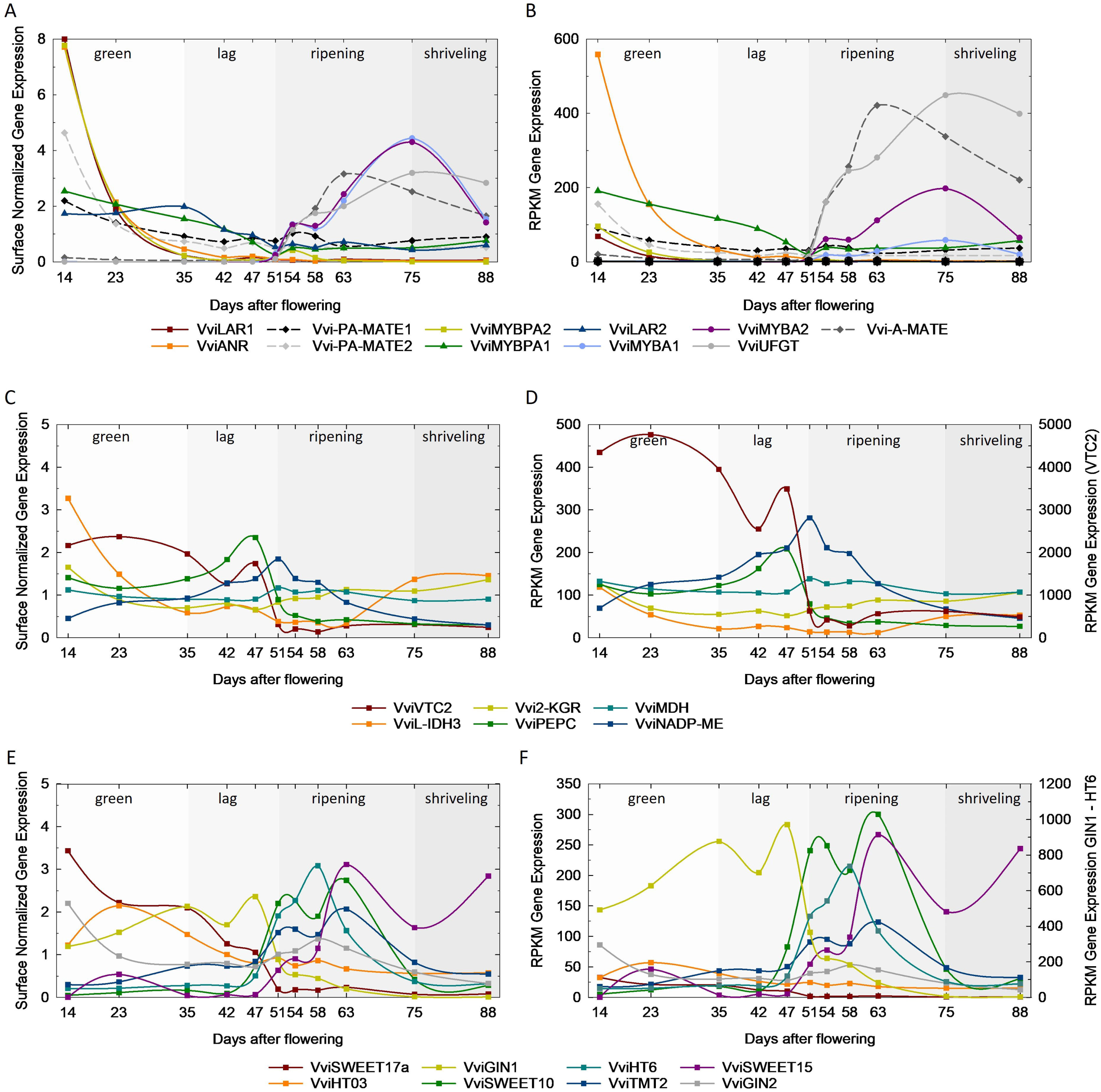
Key genes involved in proanthocyanidins and osmolytes biosynthesis and transport. *VviLAR1*, *VviANR*, and *VviMYBPA2* genes of the flavonol pathway are expressed at the very early phase of berry development, followed by *VviLAR2* and *VviMYBPA1* in the green plateau; anthocyanins-related genes as *VviMYBA1*, *VviMYBA2*, and *VviUFGT* are activated three days after berry softening (**A, B**). *VviVTC2*, *VviL-IDH*, and *Vvi2-KGR* are known biosynthetic genes of tartaric acid, while *VviPEPC*, *VviMDH*, and *VviNADP-ME* are committed to malic acid synthesis and breakdown (**C, D**). ‘Green’ sugar transporters such as *VviSWEET17a*, *VviHT3*, and *VviGIN1* are replaced at softening by *VviSWEET10*, VviHT6, and *VviTMT2*. All these genes are stopped at the arrest of phloem at 75 DAF, except for *VviSWEET15*, which remained active during shriveling (**E, F**). Graphs are expressed as surface normalized data (A, C, E) and Reads Per Kilobase per Million mapped reads (RPKM) (B, D, F). In the surface normalized graphs, each expression value was divided by the average on the 11 developmental points to reject large differences in expression among genes. Softening (S) started at an unknown date between 47 and 51 days.

Three phosphoenolpyruvate carboxylases isogenes (*VviPEPC*), committed to the synthesis of the malate precursor oxaloacetate, are expressed in berries, with *Vitvi12g00185,* the most expressed one, belonging to the black module, correlated with the rate of malic accumulation. This gene was induced at 23 DAF and peaked at the end of the lag phase at 47 DAF before decreasing during ripening when the net flux of malate turns negative. By contrast, cytosolic malate dehydrogenases (*cytMDH*, *Vitvi07g00599*, *Vitvi07g00600*) showed a constitutive expression. The mitochondrial dicarboxylate/tricarboxylate carrier (*Vitvi08g01801*) was also found in the black module, suggesting a possible involvement of mitochondrial MDH in the reduction of oxaloacetate to malate in the green stage. However, an NADP-dependent malic enzyme (*VviNADP-ME*, *Vitvi11g00272*, yellow module) (Franke and Adams, 1995; Or *et al*., 2000) deputed in the degradation pathway showed a pyramidal pattern, peaking with malic acid at the onset of ripening (51 DAF), when acidity switched from accumulation to degradation. The remanence of this transcript over the complete berry cycle is in line with the vital necessity of coping with cytosolic acidosis by immediately degrading harmful malic acid during any transient stress, causing an H^+^ leakage from the vacuole.

### Ripening guided by membrane compartmentation

Although sugar import as an energy and carbon source would principally occur through the symplastic pathway during the green phase (Zhang *et al*., 2006), different sugar transporters were specifically expressed, such as the bidirectional sugar transporter *VviSWEET17a* (*Vitvi05g00013*), the sucrose transporter *VviSUC27* (*Vitvi18g01315*), and the hexose transporter *VviHT3* (*Vitvi11g00611*). These isoforms, included in brown and black modules, were then turned off at softening. Among the sucrose metabolism genes, a sucrose synthase (*Vitvi11g00030*, *VviSuSy4*, green module) and a vacuolar invertase (*Vitvi16g00713*, *VviGIN1* black module) were specifically expressed in the green phase before decreasing during the lag phase. These genes were then suddenly replaced by the plasma membrane *VviSWEET10* (*Vitvi17g00070*), the tonoplastic *VviHT6* (*Vitvi18g00056*), and *VviTMT2* (*Vitvi03g00247*), marking the induction of the apoplasmic pathway (Fig. 5E, 5F). All these genes from the yellow module turned up at (or slightly before) softening and stopped at the phloem arrest (Savoi *et al*., 2021). *VviSWEET15* (*Vitvi01g01719*, magenta module) was induced later and persisted during late ripening. A second sucrose synthase (*Vitvi07g00353*, *VviSuSy3*, magenta module) started to increase at softening together with three sucrose phosphate synthases (VviSPS, *Vitvi11g00542*, *Vitvi05g01193*, *Vitvi18g02365*) from the magenta and yellow modules. *VviGIN2* (*Vitvi02g00512*) was less expressed than *GIN1* in the early phase of green development and even decreased in the lag phase, showing a second peak of expression during ripening.

### Proton pumps driving vacuolar hyperacidification and sugar loading

The three grapevine isoforms of V-PPase showed different expression patterns (Fig. 6A, 6B), with *VviVPP1* (*Vitvi14g00101)* showing a rather constitutive expression profile, highly expressed since the beginning of organic acid accumulation, then exhibiting a transient increase simultaneous with the peak in *VviHT6* expression and malate breakdown, before decreasing at 63 DAF while staying highly expressed until over-ripening. After having been expressed threefold less than *VviPP1* during the green stage, the H^+^PPase isoform *VviPP2* (*Vitvi11g00560,* yellow module) suddenly reached a comparable level of about 130 RPKM from the onset to the end of the sugar accumulation process, collapsing after that. Lastly, the third isoform, *VviPP3* (Vitvi09g00693), was quite constant during the first growth phase, being at approximately 50 RPKM, for suddenly peaking at 47 DAF, a few days before softening, anticipating *VviPP2*. During the ripening phase, *VviPP3* increased again at 58 DAF. The catalytic subunit A in the cytoplasmic domain V1 of V-ATPase *VviVHA-A* (*Vitvi09g01397,* yellow module) and the P-ATPase *VviPH5* (*Vitvi09g00006,* grey module), already expressed in the green stage, also increased in expression during the period of sugar accumulation starting from softening, until declining when phloem unloading and berry expansion stopped. *VviPP2* was more expressed than *VviVHA-A*, followed by *VviPH5,* and remarkably, the RPKM sum of H^+^ pumps on the tonoplast approached the huge expression of *VviHT6*. In comparison, the expression of *VviPH1* (*Vitvi07g02600*), whose interaction with *VviPH5* is critical for acidification in lemon, appeared negligible.

**Figure 6.**
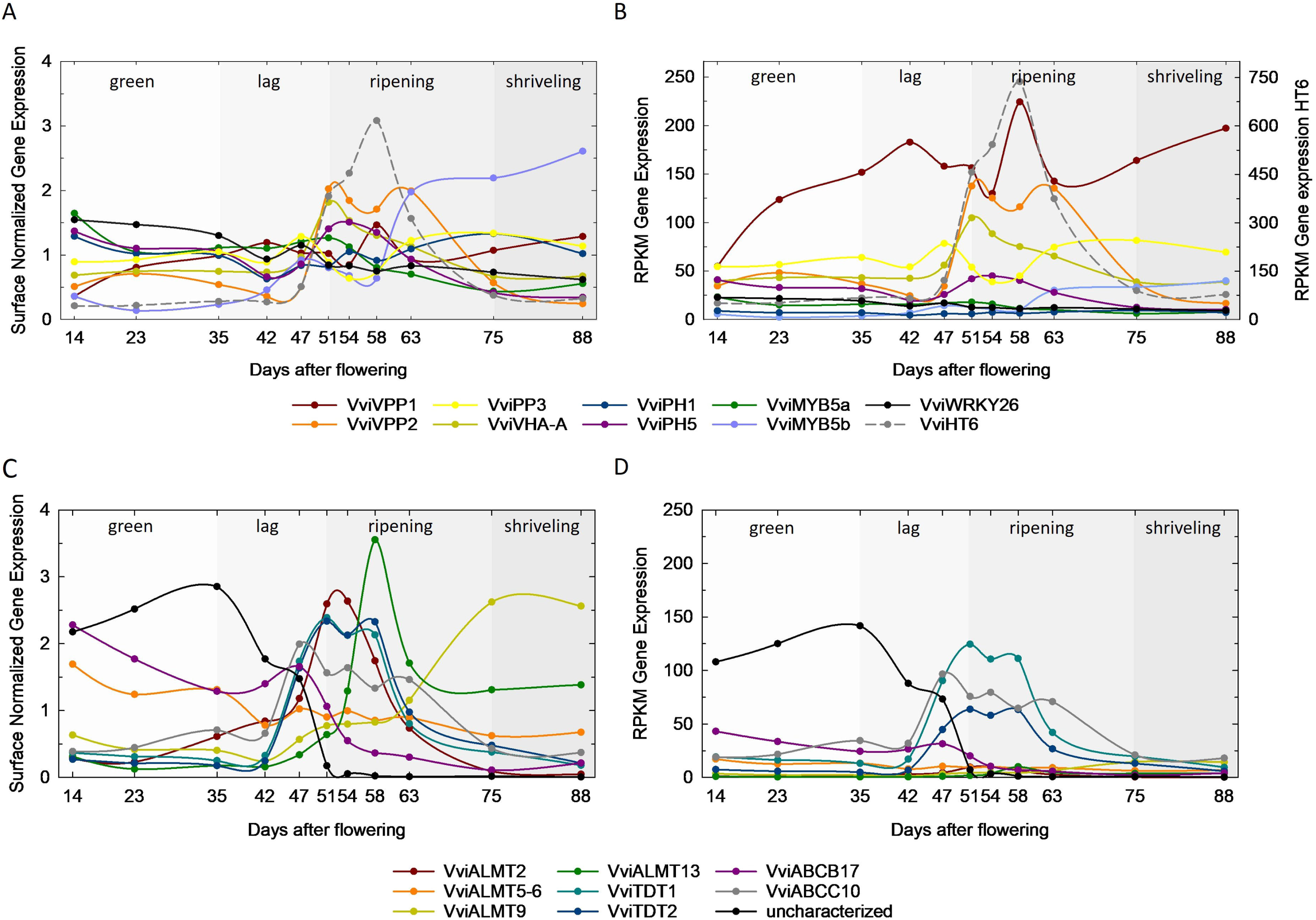
Proton pumps and transporters genes as developmental switches. The three types of tonoplastic proton pumps (V-PPase, V-ATPase, P-ATPase) displayed specific expression patterns during berry development. *VviPP2*, *VviVHA-A*, and *VviPH5* dramatically increased in expression with *VviHT6* at softening, 51 DAF. The peak of *VviPP1* mimicked the one of *VviHT6* during ripening (**A, B**). Only a few transporters were expressed in the green phase such as the *VviABCB17*, *VviALMT5-6*, and a still uncharacterized vacuolar membrane protein (**C, D**). Several ALMTs and TDT channels increased slightly before the ripening phases and the above mentioned changes in *VviHT6* and H^+^ pumps. Graphs are expressed as surface normalized data (A,C) and Reads Per Kilobase per Million mapped reads (RPKM) (B,D). In the surface normalized graphs, each expression value was divided by the average on the 11 developmental points to reject large differences in expression among genes. Softening (S) started at an unknown date between 47 and 51 days.

### Vacuolar ion transport

Globally, a set of tonoplastic ion channels and transporters (mostly anions) reached maximal expression at 51 DAF (softening, onset of ripening). The first ones, encoding for tonoplast dicarboxylate transporters (*VviTDT*, *Vitvi10g00114*, *Vitvi10g00204,* yellow module), ATP-binding cassette *VviABCC10* (*Vitvi15g00424,* yellow module) and *VviABCB17* (*Vitvi08g01627,* tan module) were induced one week before softening (DAF 47). The expression pattern of VviTDTs was quite parallel to the rate of malate breakdown. They were followed by *VviALMT2* (*Vitvi06g00922*), then *VviALMT13* (*Vitvi02g00066*), and only during late ripening by *VviALMT9* (*Vitvi17g00333,* blue module). The only transporter (*Vv*i*ALMT5-6* - *Vitvi01g00266*), which seemed to be faintly expressed in the green period (belonging to the green module), was low expressed, questioning its central role in the entry of organic acid during the accumulation period. The only tonoplastic membrane protein (*Vitvi09g00350*, brown module) whose expression precisely aligns with the malate accumulation kinetics in immature green berries is still uncharacterized in plants, being highly expressed in the green phase for then declining at the onset of ripening and staying off during the ripening phase (Fig. 6C, 6D).

### Ripening hormone signaling and transcription factors

Modules pink and yellow provide precise insights into the molecular cascade triggering the onset of ripening since the first one displayed the highest association with the lag phase (0.73) and, in particular, the stage L5 (0.63), just four days before the second one, highly correlated with the softening phase (0.77). The expression of 1-aminocyclopropane-1-carboxylate oxidase (*VviACO1* - *Vitvi10g02409*, pink module), committed to ethylene biosynthesis, started to increase at 23 DAF and peaked during the lag phase at 47 DAF, four days before the onset of ripening. It sharply decreased after that, becoming barely detectable from 58 DAF onwards (Fig. 7A, 7B). The expression of the downstream signaling gene, ethylene insensitive 3 (*VviEIN3* - *Vitvi13g01126*), also increased in the green stage, peaking at softening (51 DAF). However, *VviEIN3* showed a second and slightly highest peak of expression at 75 DAF during late ripening stages, as previously documented (Cramer *et al*., 2014). Nevertheless, the peak of ethylene-related genes preceded the ABA-related ones, and the decrease of *VviACO1* was relayed by the induction of *VviNCED2 - Vitvi10g00821*. However, *VViNCED3* - *Vitvi19g0135*, silent during the green phase, was largely induced at 47 days, although reaching its maximal expression at 54 DAF, three days after softening. It therefore appears as the first abrupt signal announcing softening. Notably, the ABA-responsive transcription factor *VvibZIP45,* also known as *VviABF2* (*Vitvi18g00784*, yellow module), showed a very close pattern with *VviNCED3* (Fig. 7A, 7B). *VviNCED2*, which was two times less expressed than its homolog, suddenly increased its expression at softening (51 days), peaking simultaneously with *VViNCED3* at 54 days, then decreased until being switched off simultaneously with phloem unloading.

**Figure 7.**
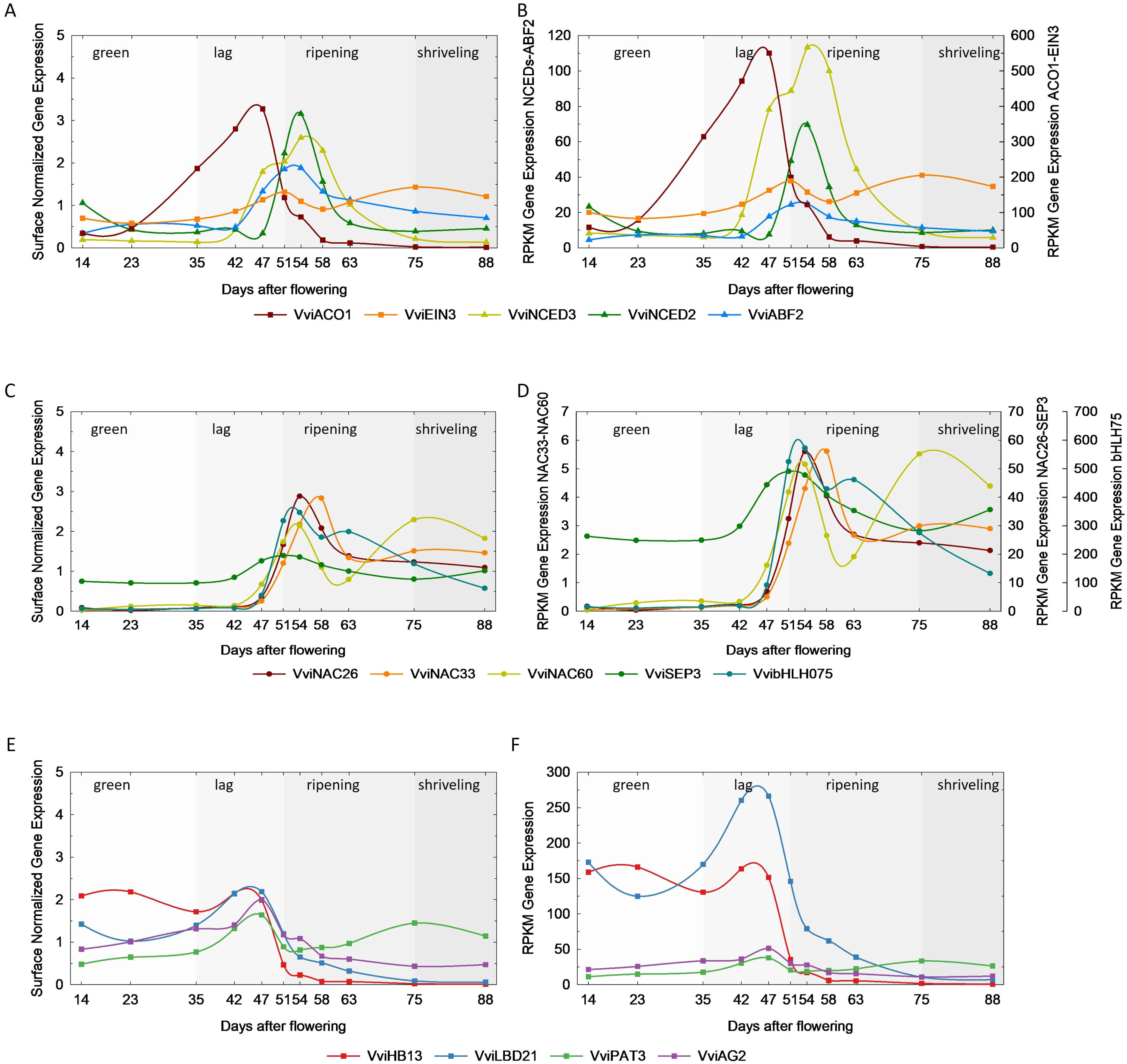
Fruit hormones and molecular markers of the onset of ripening. Due to the induction of *VViNCED3*, the *VviACO1/VViNCED3* (or Ethylene/ABA) balance was already impaired four days before softening, announcing the burst in *VViNCED2* expression (**A, B**). Transcription factors, such as *VviNAC26*, *VviNAC33*, *VviNAC60*, *VviSEP3*, and *VvibHLH075*, already detected at 47 days, show an obviously enhanced expression at the onset of ripening in line with their possible control of transcriptome reprogramming at this stage (**C, D**). Other putative transcription factors (*VviHB13*, *VviLBD21*, *VviPAT3*, *VviAG2*) may negatively control ripening, being enhanced during the lag phase and abruptly turned off at softening (**E, F**). Graphs are expressed as surface normalized data (A,C) and Reads Per Kilobase per Million mapped reads (RPKM) (B,D). In the surface normalized graphs, each expression value was divided by the average on the 11 developmental points to reject large differences in expression among genes. Softening (S) started at an unknown date between 47 and 51 days.

Based on literature data, WGCNA module membership, and gene expression profiles refined by cluster analysis (Savoi *et al*., 2023), we identified a series of transcription factors showing diverse expression ranges when peaking at S7, and thus possibly involved in the fruit ripening program (Fig. 7C, 7D). This included NAC members such as *VviNAC60 - Vitvi08g01843* (D’Incà *et al*., 2023), *VviNAC26 - Vitvi01g01038* (Zhang *et al*., 2021), *VviNAC33 - Vitvi19g01484* (D’Incà *et al*., 2021), a basic helix-loop-helix *VvibHLH075 - Vitvi17g00046* (Fasoli *et al*., 2018), and *VviSEP3 - Vitvi01g01677* from the MADS-box family (Mellway and Lund, 2013). However, anion transporters and hormones previously discussed prone us to look for signals activating the transcriptome reprogramming at L5, four days before the phenological softening stage (Fig. 7E, 7F). This could identify a homeodomain leucine zipper *VviHB13* - *Vitvi01g00958* (Terrier *et al*., 2005), a lateral organ boundaries domain gene of class I c3 *VviLDB21* - *Vitvi13g00085* (Grimplet *et al*., 2017), a GRAS transcription factor *PAT3* - *Vitvi10g00271* (Grimplet *et al*., 2016), and finally the MADS-box AGAMOUS 2, *VviAG2* - *Vitvi10g00663* (Mellway and Lund, 2013; Green *et al*., 2024).

## Discussion

For the first time, developmental changes in the net fluxes of water, hexose, malic and tartaric acids were estimated and correlated to gene expression at their pertinent single fruit regulatory level. Image analysis allowed us to quantify the double sigmoidal growth curves non-destructively, before sampling berries with very high time resolution, considering their own individualized internal clock rather than the observer one. Getting rid of the developmental chimerism inherent to averaging asynchronous berries (Bigard *et al*., 2019; Shahood *et al*., 2020) allowed us to unravel metabolically and transcriptionally homogeneous phases, whose durations were shortened by two weeks at least, leading to a noticeable acceleration of metabolic rates.

### Phenological transitions are faster at the onset of ripening

Transitions between physiological or transcriptomic stages appeared extremely abrupt on single berries, allowing us to subdivide berry development into more accurate phases than previously considered. For instance, significant transcriptomic changes occurred between S6 and S7, which were collected three to four days apart after their individual softening dates. When the incipit of softening is unknown, they remain indistinguishable in the ‘soft green’ class, so their transcriptomes would be averaged in phenotypically sorted or stratified samples (Hernández-Montes *et al*., 2021). The outstanding transcriptomic homogeneity of single, duly sorted berries inside triplicates indicates that the comparatively large noise observed on average samples (i.e., Tornielli *et al*., 2023) reflects random variations in their age pyramid. Consequently, single-berry transcriptomes led the two first PCA components to explain a higher degree of variance in gene expression (54% and 15%, respectively, Fig. 2B) without eliminating any noisy gene. Although some divergence appeared among triplicates after the R9 stage, individual transcriptomes did not depart from the 2D plot, excluding any microenvironmental scattering effect and simply indicating that the transcriptomic clock is more accurate than our preliminary synchronization procedure. Actually, R9 berries were sorted by considering a 50% volume increase since softening. Present results suggest that, although limited in the 20 berries whose complete developmental cycle was monitored here, some variation exists in berry expansion at the same developmental stage, as recently indicated by high throughput automated monitoring of berry growth (Daviet *et al*., 2023). After maximum berry volume, when the berry turns hydraulically disconnected from the vine (Savoi *et al*., 2021) (Fig. 4A, 4B), the transcriptome turns quite stable: R10 and Sh11 samples are grouped (Fig. 2B), as the transcriptomic time decelerated (Fig. 2C).

The spatiotemporal separation in the synthesis of specific phenolic compounds is well documented during grape berry development (Czemmel *et al*., 2012). Proanthocyanidin biosynthesis is modulated by *VviMYBPA1* and *VviMYBPA2* (Bogs *et al*., 2007; Terrier *et al*., 2009). Single berry transcriptomics allows to resolve the expression patterns of these TF just separated a few days apart. In this respect, *VviMYBPA2* strongly decreased before 23 DAF, simultaneously with the core PA gene *VviLAR1* and *VviANR.* Moreover, among other water channels isogenes, *VviPIP2.7* and *VviTIP2.1* were predominantly expressed during the first growth phase, when water would be principally translocated by xylem and the limited phloem unloading essentially symplastic, thereby excluding any significant transmembrane fluxes in sieve tubes (Fig. 4B). *VviPIP2.7* was replaced by *VviPIP2.5* and *VviPIP2.3*, when shifting to predominating apoplastic phloem unloading, being these isoforms most probably sieve element-specific (Stanfield *et al*., 2017). Conversely, *VviPIP1.3* showed a comparable and constitutive expression from 23 DAF until the phloem stopped, possibly ensuring continuous water movement inside the flesh. *VviTIP1.2*, reaching during ripening the same RPKM as *VviTIP2.1* during the green stage, may thus also appear ‘flesh specific’. The adaptive mechanisms below this shift in TIP isogenes is questionable. Finally, RPKM of expansin genes indicated a higher expression during ripening of *VviEXP14* and *VviEXP19* compared to other isogene expressed in green phase, Fig. 4D, (Malacarne *et al*., 2024). The two successive waves of expansin activated during ripening may respectively correspond to softening and growth resumption.

### Sequence of ripening events

Kinetic analysis of individual berries confirms that the first ripening phenotype indicators are a drop in fruit firmness and a reduction in the G/F ratio (Fig. 1C). These changes co-occurred with a pronounced surge in the intake and storage of photoassimilates above 0.15 M hexoses, as previously documented (Bigard *et al*., 2022). *VviSWEET10* and *VviHT6* sugar transporters induced on the plasma and tonoplast membranes between 47 and 51 DAF announced this huge acceleration of phloem unloading. Berries analyzed through continuous image analysis softened at 51 ± 1.5 DAF, with a notable 7 ± 1.5 days delay before growth resumption. This early ripening sequence agrees with previous reports on which growth was lacking (Gouthu and Deluc, 2015; Castellarin *et al*., 2016; Hernández-Montes *et al*., 2021) and confirms that berry expansion would not resume before 0.4 M hexoses (Shahood *et al*., 2020). Clearly, single berry monitoring allows distinguishing developmental events previously thought to be simultaneous in all studies based on average samples and considering 50% of colored berries (mid-veraison) as a rough indicator of the onset of ripening (i.e., Fasoli *et al*., 2018; Theine *et al*., 2021). In fact, both *VviMYBA1*, *VviMYBA2* TFs and the regulatory gene *VviUFGT* simultaneously enhanced their expression at 54 DAF, four days after softening (Fig. 5A, 5B). Recently, the first high throughput automated analysis of time-lapse images led to the conclusion that there is a 4.4-day average delay between growth resumption and coloration in the *Vitis vinifera* cv Alexandroouli (Daviet *et al*., 2023). However, one should not reject that this delay should depend on genetic differences and environmental conditions like light, water, or crop load. Merging growth and sugar concentration data evidenced a progressive acceleration in hexose accumulation at softening, finally reaching 64 µmol hexose*min-1 N berry-1. This net accumulation rate is four times faster than can be calculated on average, unsynchronized berry samples (i.e., 16 µmol hexose*min-1 N berry-1 for a 6-week period in Fasoli *et al*., (2018)). Malic acid breakdown started at approximately 0.3 M sugar (Fig. 1D), and the initial rate of malate breakdown corresponds to a 1:1 exchange of sucrose for H^+^ at the tonoplast (Fig. S1), as recently shown on different varieties (Shahood *et al*., 2020; Bigard *et al*., 2022).

Fruit ripening involves various hormonal signals and interactions with several TFs. As a non-climacteric fruit, ripening relies on a composite interplay among hormones at the onset of ripening, among which the ABA/auxin balance and ethylene (Zenoni *et al*., 2023). Increased time resolution gained on single berry put forwards *VviNCED3*, the main ABA biosynthetic gene, as a first abrupt signal preceding the drop in the ethylene *VviACO1* and the induction of *VviNCED2* at softening (Fig. 7B). Finally, several transcription factors already characterized in the literature appeared clearly induced shortly before softening (Fig. 7E, 7F) or peaking a few days later at S7 (Fig. 7C, 7D). Further studies are needed to establish their respective roles in fruit ripening.

### Unraveling the central role of vacuolar energetics on the sugar acid nexus

The net sugar accumulation and malate breakdown rates are consistent with the sudden induction of global sucrose/H^+^ exchange at the onset of ripening when the *VviHT6/VviSWEET10* transcripts are switched on. The rate of sugar accumulation progressively accelerates like the expression of *VviSWEET15*; however, unlike the previous ones, this last gene is not switched off at the completion of sugar loading. Sugar storage would be first energized by the vacuolar acidity gradient previously formed in the green stage upon malic acid accumulation, the sugar/H^+^ exchange being charge compensated by the release of vacuolar malate, triggering its mitochondrial oxidation. However, sugar loading accelerates while the malic acid content vanishes (Fig. S1). Transport experiments on tonoplast vesicles suggested that H^+^ pumps progressively take over the electro-neutralization process (Terrier *et al*., 2001). The simultaneous increase of *VviHT6* and vacuolar H^+^ pump transcripts (Fig. 6A, 6B) suggests that a phenotypically silent H^+^ recirculation could parallel or relay malate efflux. This could indicate that (1) hexose, not sucrose, would be transported by *VviHT6*, (2) a fail-safe mechanism preventing excessive release of acidity when compared to oxidative capacity must be anticipated to provide instant response upon stress, or (3) due to a lack of selectivity of the malate channel, non-metabolizable tartrate is released in the cytoplasm, requiring an active return to the vacuole. Whatever, with a respiratory rate of about 12-20 µmolO_2_min-1Nberry-1, hence a maximum of 110 µmol ATP min-1Nberry-1 (recalculated from Morales *et al*., 2024), the intensity of sugar loading in individual berries turns hardly compatible with the energetics of a classical import pathway involving cell wall sucrose inversion and hexose H^+^ symporter at the plasma membrane, which would consume circa 55% of ATP possibly formed by oxidative phosphorylation. This energy-guzzling pathway is thus bypassed by *VviSWEET10* expression, which closely matches the one of *VviHT6*, preceding the one of *VviSWEET15*.

### Tonoplastic proton pumps are simultaneously expressed

Within fruit cells, malate primarily undergoes synthesis in the cytoplasm, a process achieved by PEPC, MDH, and possibly malate synthase (Sweetman *et al*., 2009; Etienne *et al*., 2013). Its synthesis from hexose also produces two harmful H^+^ retro-controlling PEPcase and activating ME, so H^+^ transport from the cytoplasm into vacuoles pulls the overall malate synthesis and accumulation process. Here, we confirm that the three types of H^+^ pumps are simultaneously expressed on the tonoplast (V-PPases, V-ATPases, and P-ATPases) in green and ripe berries. The co-expression of these pumps in the green stage apparently contradicts the thermodynamic need to replace V-PPases and V-ATPase with a more electrogenic P-ATPase complex evoked on citrus (Strazzer *et al*., 2019). As a matter of fact, the *in vivo* ΔG_PPi_ of -27.3 kJ/mol (Davies *et al*., 1993) allows a 1H^+^/PPi PPiase to form a 4.7 pH units gradient across the tonoplast, and a similar conclusion holds for a 2H^+^/ATP vATPase fed with ΔG_ATP_ of -54.6 kJ/mol. These conservative values (nmin:1.75 H^+^/ATP; max oxphos ΔG_MgATP_:-61+/-3 kJ/mol) (Roberts *et al*., 1985; Davies *et al*., 1994; Wiseman *et al*., 2023) indicate there is no strict requirement for more electrogenic H^+^ pump in green berries at pHv=2.7. PH5, erroneously considered a contaminant in Terrier *et al*. (1998), is also co-expressed with V-ATPase in apples and pears and promotes malate accumulation in these fruits (Song *et al*., 2022; Huang *et al*., 2023). The presence of two ATPases with possibly different H/ATP ratios is puzzling, so further work is needed to establish whether these pumps operate in parallel on the same membrane.

### Open questions on malate entry and exit route

Questions remain on malate’s entry and exit route to and from the vacuole. Unexpectedly, the expression of *bona fide* malate/tartrate vacuolar transporters did not parallel the vacuolar influx of organic acids during the green stage (Fig. 6C 6D), neither that of many other anion transporters. The only aluminum-activated malate transporter family member characterized in grapevine is *VviALMT9* on chromosome 17. This gene has been pointed out to mediate inward-rectifying malate and tartrate currents across the tonoplast (De Angeli *et al*., 2013). However, its expression during ripening does not match the timing of malate entry in the green phase (Fig. 6C 6D). ALMT phylogenetic analysis (Dreyer *et al*., 2012; Rienth *et al*., 2016) indicated that additional clade II genes should be targeted to berry tonoplast, such as the *VviALMT5-6*, misannotated as a unique gene in the grapevine chromosome 1. Similarly, on apple chromosome 16, the Ma locus comprises two genes, *MDP244294* and *MDP252114* (Bai *et al*., 2012), sharing the same conserved duplicated structure as in *V. vinifera* chromosome 1, and not on chromosome 17. Therefore, alignment and synteny confirm *VviALMT5-6* as the true orthologs of the Ma locus, controlling the genetic diversity of malic acid in apple. Actually, these ALMT were constitutively but slightly expressed during pericarp development, prompting the question of the effective player of malate vacuolar import during the green stage. In this respect, by digging the present dataset, we identified a new putative anion channel still uncharacterized in plants (*Vitvi09g00350, D7U0H3*) whose expression perfectly matched the malate accumulation kinetics in green berries. Although uncommented, such expression profile is confirmed in other transcriptomic data (i.e., Rienth *et al*., 2016; Fasoli *et al*., 2018). Finally, the transporter was localized on the vacuole during a pioneering proteomic study and highly abundant during the green phase (Kuang *et al*., 2019). Further studies are needed to characterize the functional role of this gene.

## Conclusion

Our work provides a resource on single-fruit transcriptomes and paves the way for comparative analyses of acidic fleshy fruit development in different perennial species, both non-climacteric and beyond. Time-resolved changes in gene expression highlighted the multi-faceted adaptation of membrane transports retained by evolution, allowing the pericarp to rapidly import sugar, before signaling this nutritional reward with anthocyanin. Announced by concerted changes in hormonal, kinetic, and membrane transporters a few days before softening, the sudden induction of sugar accumulation and malate breakdown coincides with the activation of vacuolar transport systems (*VviHT6* and *VviSWEET10*) at the onset of ripening. The findings suggest a model where the vacuolar acidity gradient drives sugar storage while triggering the malate oxidation processes and the actuation of H^+^ pumps. Finally, known ALMT transporters, like *VviALMT9*, do not align with malate accumulation in the green phase. Instead, an uncharacterized anion transporter (*Vitvi09g00350*) was strongly expressed, hinting at its potential role in vacuolar malate import. Further functional studies are needed to clarify the role of specific transporters in malate movement during berry development.

## Supplementary materials

Figure S1. The stoichiometry of malic acid breakdown versus sugar accumulation.

Table S1. List of samples, their acronym, and phenological stages of reference.

Table S2. Parameters of the sigmoidal function calculation for the 20 reference berries.

Table S3. Reconstructed curve of berry growth

Table S4. Metabolite fluxes in every berries sampled from anthesis to overripening.

Table S5. Top-100 PC1 and PC2 positive and negative loadings.

Table S6. Transcriptomics time.

Table S7. WGCNA gene module association and membership correlation.

## Acknowledgments

We thank the Department of Viticulture and Enology of Institut Agro Montpellier for facilitating access to the experimental vineyard Pierre Galet and the genomic platform of INRAE managed by Dr. Sylvain Santoni for helping in sample preparation and processing.

## Author contributions

LT and CR: conceptualization; SS, LT, and CR: methodology; SS, MS, GS, AW: formal analysis; SS, LT, and CR: investigation; SS, LT, and CR: resources; SS and CR: data curation; SS: writing - original draft; SS, LT, and CR: writing - review & editing; LT: funding acquisition.

## Data Availability

All raw transcriptomic reads have been deposited in the NCBI Sequence Read Archive (http://www.ncbi.nlm.nih.gov/sra). The BioProject is PRJNA862686.

